# Photoaffinity labeling of protein targets in a complex metazoan: proof-of-concept using a probe for *Schistosoma mansoni* tubulin

**DOI:** 10.1101/2025.10.04.680471

**Authors:** Bobby Lucero, Thibault Alle, Marianna Bufano, Yuemang Yao, Kurt R. Brunden, Andrea Brancale, Carlo Ballatore, Karol R. Francisco, Conor R. Caffrey

## Abstract

Treatment of schistosomiasis, a prevalent neglected tropical disease, relies precariously on a single drug. The discovery and development of alternative anti-schistosomal small molecules most often relies on phenotypic (whole-organism) screening, whereas target- or “protein-first”-based discovery options are hampered by a paucity of genetic interrogation strategies and a complex biology. Here, we demonstrate the application of photoaffinity labeling (PAL) as a chemical biology strategy to probe protein target-ligand interactions in the schistosome. Using a triazolopyrimidine (TPD) probe that binds tubulin, we established a PAL workflow with living *Schistosoma mansoni* worms. The probe elicited deleterious phenotypic responses consistent with the TPD series and, upon UV light activation, covalently labeled tubulin as identified by proteomics. Specific concentration-dependent engagement of tubulin was confirmed using a photostable competitor TPD. When applied directly to worm lysates, the PAL workflow produced non-specific labeling, suggesting that the conformation of the protein target is important for ligand binding. The successful application of PAL for a metazoan is, to our knowledge, novel, and the platform should prove generally applicable to identifying potential drug targets, and exploring protein-ligand interactions, in schistosomes and other organisms.

## MAIN TEXT

Schistosomiasis, a neglected tropical disease (NTD) caused by parasitic flatworms of the genus *Schistosoma*, affects approximately 700 million people worldwide,^1, 2^ with over 250 million requiring chemotherapy.^3^ This chronic infection imposes a substantial health and socioeconomic burden in endemic regions, contributing to long-term morbidity and reduced productivity.^4, 5^ Notwithstanding its global impact, treatment relies on just one drug, praziquantel (PZQ).^6, 7^ Although safe and reasonably effective (50–90%^8, 9^ cure), it has notable limitations, particularly its lack of activity against juvenile worms^10, 11^ which allows infection and associated morbidity to continue as the parasites mature and lay eggs. In addition, PZQ displays suboptimal pharmacokinetics, characterized by extensive first-pass metabolism to therapeutically inactive products.^12, 13^ Finally, the reliance on PZQ heightens the risk of drug resistance.^14^ Collectively, these factors underline the need for new therapeutics, including those with novel mechanisms of action.

The discovery and structural-activity characterization of anti-schistosomal small molecules has most often employed phenotypic (whole-organism) screening of the pathogen in culture.^15-17^ Target- or ‘protein-first’ based discovery methods are less common and have been hindered by both biological and technical constraints.^18^ For example, many of the reverse and forward genetics technologies to investigate gene product function in mammalian cells,^19, 20^ some protozoan parasites,^21, 22^ and the free-living nematode, *C. elegans*.^23^ are not yet routine for the schistosome, in spite of considerable effort.^24-26^ An exception is transient (non-germline) RNA interference which has demonstrated robust utility over the last 25 years to identify gene products that influence parasite growth and/or survival.^27-32^ Added to these difficulties are the more general constraints regarding the pathogen’s complex life-cycle, involving invertebrate and vertebrate hosts, and the need for animal biosafety level (ABSL)-2 infrastructure.

In this context, we investigated a chemical biology approach to identify drug targets in *S. mansoni*. Specifically, we identified small molecule pyrimidines that engage microtubules (MTs) in mammalian cells and elicit deleterious phenotypic responses in *ex vivo* adult *Schistosoma mansoni* worms.^33^ We subsequently repurposed these compounds against another pathogen, the protozoan, *Trypanosoma brucei*,^34^ and designed and synthesized a photoaffinity labeling (PAL) probe (**1**, Table 1) that labeled *T. brucei* tubulin.^35^ Inspired by these findings, we hypothesized that a PAL strategy could be applied to *S. mansoni* to identify molecular targets under conditions similar to those used in our phenotypic assays. Accordingly, we developed a protocol using the validated tubulin-targeting probe, **1**,^35^ to successfully label schistosome tubulin in a concentration-dependent manner. The data demonstrate that PAL can be effectively combined with phenotypic screens to investigate mechanisms of action, thus providing an exciting opportunity to identify protein targets in an experimentally challenging organism.

**Table 1.**
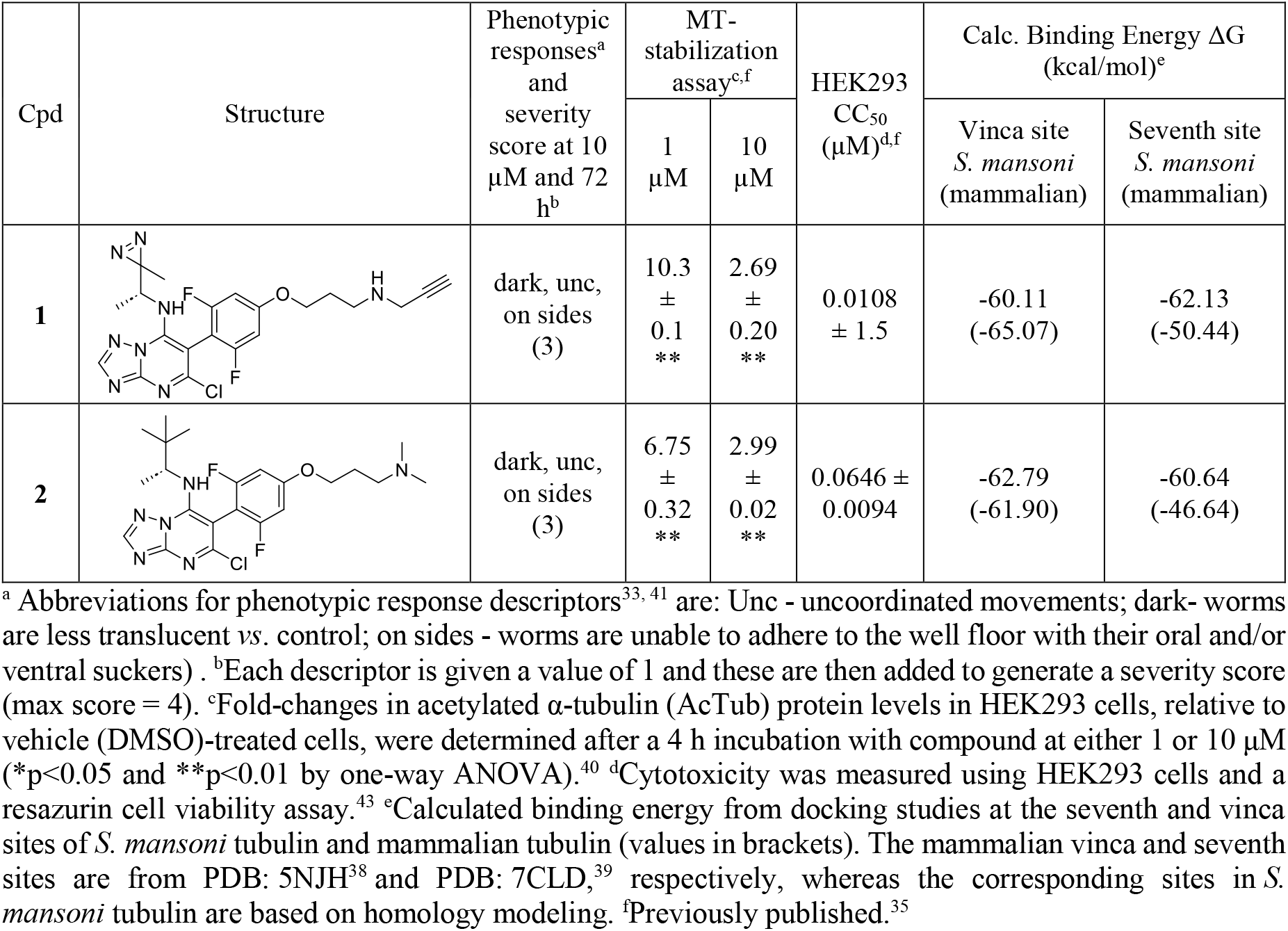
Bioactivity of PAL probe **1** and photostable competitor **2** *vs*. adult *S. mansoni*, MT-stabilizing Activity (AcTub), cytotoxicity in HEK293 cells, and binding energy ΔG values based on molecular docking.

| Cpd | Structure | Phenotypic responses<sup>a</sup> and severity score at 10 μM and 72 h<sup>b</sup> | MT-stabilization assay<sup>c,f</sup> | | HEK293 CC<sub>50</sub> (μM)<sup>d,f</sup> | Calc. Binding Energy $\Delta G$ (kcal/mol)<sup>e</sup> | |
| --- | --- | --- | --- | --- | --- | --- | --- |
| | | | 1 μM | 10 μM | | Vinca site *S. mansoni* (mammalian) | Seventh site *S. mansoni* (mammalian) |
| **1** | | dark, unc, on sides (3) | 10.3 ± 0.1 ** | 2.69 ± 0.20 ** | 0.0108 ± 1.5 | -60.11 (-65.07) | -62.13 (-50.44) |
| **2** | | dark, unc, on sides (3) | 6.75 ± 0.32 ** | 2.99 ± 0.02 ** | 0.0646 ± 0.0094 | -62.79 (-61.90) | -60.64 (-46.64) |

<sup>a</sup> Abbreviations for phenotypic response descriptors<sup>33, 41</sup> are: Unc - uncoordinated movements; dark- worms are less translucent vs. control; on sides - worms are unable to adhere to the well floor with their oral and/or ventral suckers). <sup>b</sup> Each descriptor is given a value of 1 and these are then added to generate a severity score (max score = 4). <sup>c</sup> Fold-changes in acetylated α-tubulin (AcTub) protein levels in HEK293 cells, relative to vehicle (DMSO)-treated cells, were determined after a 4 h incubation with compound at either 1 or 10 μM (\* $p$ <0.05 and \*\* $p$ <0.01 by one-way ANOVA).<sup>40</sup> <sup>d</sup> Cytotoxicity was measured using HEK293 cells and a resazurin cell viability assay.<sup>43</sup> <sup>e</sup> Calculated binding energy from docking studies at the seventh and vinca sites of *S. mansoni* tubulin and mammalian tubulin (values in brackets). The mammalian vinca and seventh sites are from PDB: 5NJH<sup>38</sup> and PDB: 7CLD,<sup>39</sup> respectively, whereas the corresponding sites in *S. mansoni* tubulin are based on homology modeling. <sup>f</sup> Previously published.<sup>35</sup>

Previously, we evaluated MT-active phenylpyrimidine (PPDs) and triazolopyrimidine (TPD) compounds against *ex vivo* adult *S. mansoni*.^33^ These induced various phenotypic responses that included uncoordinated movements, an inability of the oral and ventral suckers to adhere to the well and, subsequently, degeneration.^33^ Given the high amino acid sequence identity between mammalian and *S. mansoni* α- and β-tubulins, we hypothesized that these small molecules exert their effects through binding to schistosome tubulin. Due to the lack of established functional assays to directly evaluate MT-targeting activity in *S. mansoni*, we employed the TPD PAL probe, **1** (see **Table 1**), previously validated in HEK293 cells and *T. brucei*, as a chemical probe to investigate target engagement.^35^

Probe **1** was designed based on an established structure–activity relationship (SAR) for this compound class with respect to mammalian MT-stabilizing activity, specifically using TPD **2** (**Table 1**) as a structural template, and was synthesized as previously described.^34-37^ Probe **1** incorporates a diazirine group which, upon UV activation, covalently binds its protein target. An alkyne handle allows conjugation to a bifunctional reporter that comprises a TAMRA group for fluorescence imaging and biotin for affinity enrichment.^35^

In mammalian cells, TPD compounds bind to two MT sites: the vinca and the seventh sites, which are at the interface of β1-α2 and α2-β2 tubulin, respectively.^38, 39^ Molecular docking studies with probe **1** indicate comparable binding affinities for the vinca site of *S. mansoni* and mammalian tubulin, with predicted ΔG values of –60.11 and –65.06 kcal/mol, respectively (**Table 1**). Compound **2** displayed similar affinities, with ΔG values of –62.79 kcal/mol for *S. mansoni* and –61.90 kcal/mol for mammalian tubulin. Interestingly, both probe **1** and **2** exhibited greater predicted affinity for the seventh site of *S. mansoni* tubulin compared to mammalian tubulin, with ΔG values of –62.12 and –50.44 kcal/mol for **1**, and –60.64 and –46.64 kcal/mol for **2** (**Table 1**).

Examination of the docking poses of **1** and **2** with mammalian and *S. mansoni* tubulin revealed some key similarities in their binding modes (**Figure 1**). At the vinca site, for both **1** and **2**, π–π stacking is observed between the TPD core and β-Tyr222, while the dimethylamino chain is predicted to form contacts with β-Lys174, β-Val175 and β-Ser176. In contrast, at the seventh site, **1** and **2** display stronger binding affinity for *S. mansoni*, likely due to differences in interactions within surrounding residues. In both the mammalian and *S. mansoni* seventh sites, interactions include H-bonding with α-Asp211 and π–π stacking of the TPD and difluorophenyl rings with α-Tyr224 and α-Tyr210, respectively. In the mammalian seventh site, the 3,3-dimethylbutan-2-yl group is observed near α-Leu227, β-Leu246, β-Lys350 and β-Thr351, while the dimethylamino chain shows hydrophobic contacts with α-Val177 and α-Ser178. In contrast, within the *S. mansoni* seventh site, the 3,3-dimethylbutan-2-yl shows hydrophobic contacts with β-Leu246, β-Ile345, and β-Pro346, while the dimethylamino chain is surrounded by α-Gln176, α-Ile177, α-Ala178 and α-Thr179 (**Figure 1**).

**Figure 1.**
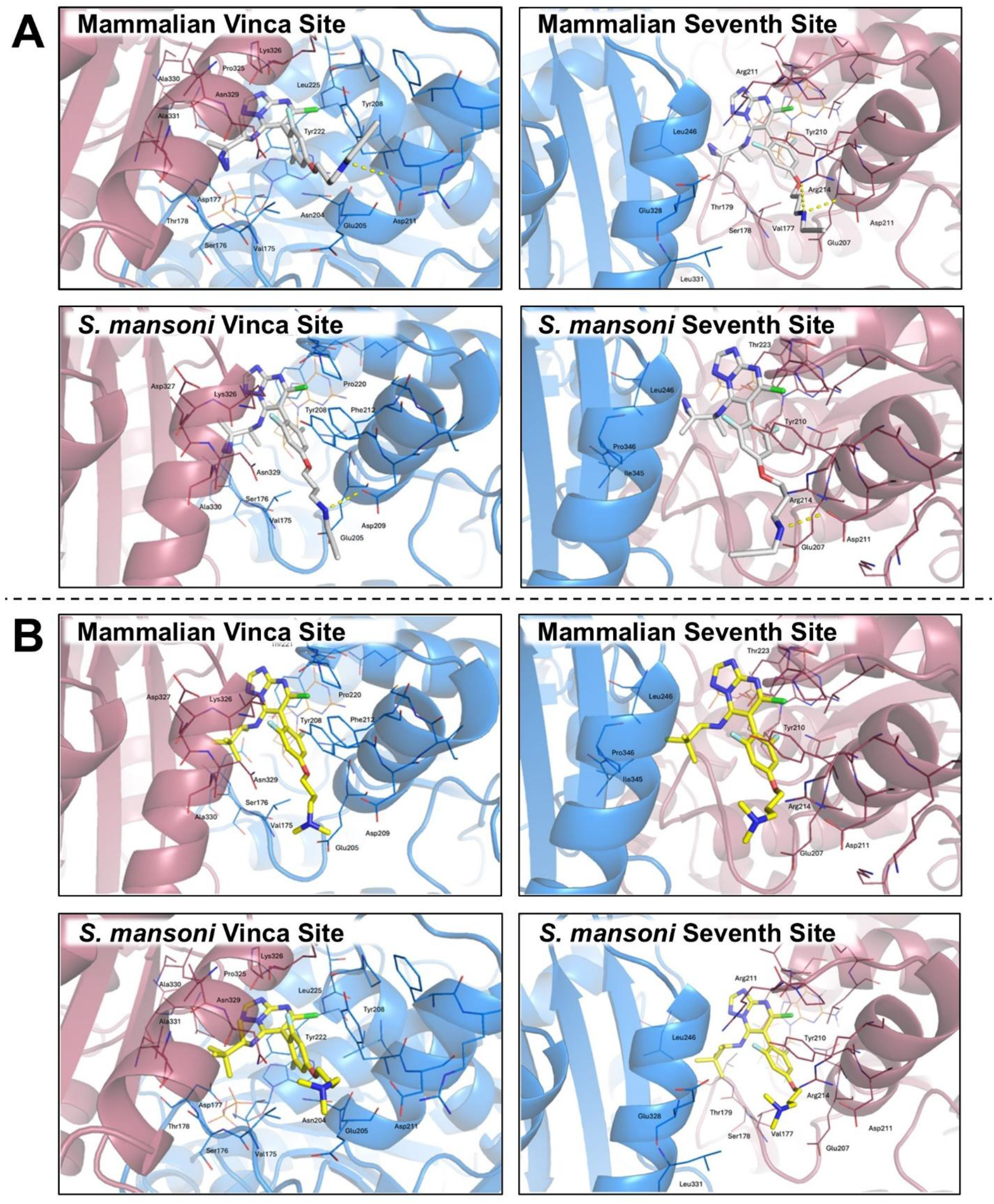
Docking of **1** (A, in grey) and **2** (B, in yellow) to the vinca (PDB: 5NJH^38^) and seventh sites (PDB: 7CLD^39^) of mammalian and *S. mansoni* tubulin (homology model): α-tubulin is shown in red and β-tubulin in blue.

Compounds **1** and **2** were evaluated for MT-stabilizing activity and cytotoxicity in HEK293 cells, as previously described (**Table 1**).^35^ Both compounds exhibited comparable MT-stabilizing activity, as evidenced by increased levels of acetylated tubulin, which is a well-established marker of stabilized MTs.^40^ The compounds were also assessed in our *ex vivo S. mansoni* phenotypic screening platform,^33, 41, 42^ which employs a constrained nomenclature of descriptors to adjudicate phenotypic changes in the parasite, and from which a severity score, on a scale of zero (no effect) to 4 (maximal effect), is calculated. After 72 h, both **1** and **2** elicited similar phenotypic responses in adult worms, characterized by darkening, uncoordinated movement, and the worm’s inability to use the oral and/or ventral sucker to adhere to the surface of the assay plate wells (**Table 1**).

Given the ability of **1** to engage MTs in mammalian cells and having recorded measurable phenotypic changes in *S. mansoni*, we developed a PAL workflow^35, 44^ (**Figure 2A**) with *ex vivo S. mansoni* adults to measure whether **1** binds to schistosome tubulin (see Supporting Information). Briefly, approximately five males and two females were incubated with 2 µM PAL probe for 2 h in a 24-well plate. Then, the probe was photoactivated for 30 min using a handheld UV lamp (365 nm) positioned 1-2 cm above the surface of the culture medium in the assay plate (minus the plate lid). Following photoactivation, worms were homogenized by mechanical sonication, and the lysate clarified by centrifugation. Proteins were subjected to a Click reaction^44, 45^ with TAMRA-biotin-azide to allow for in-gel fluorescence visualization of the labeled protein bands. This was followed by excision of the protein bands and neutravidin-based enrichment of the labelled proteins for mass spectrometry-based proteomics (see Supporting Information).

**Figure 2.**
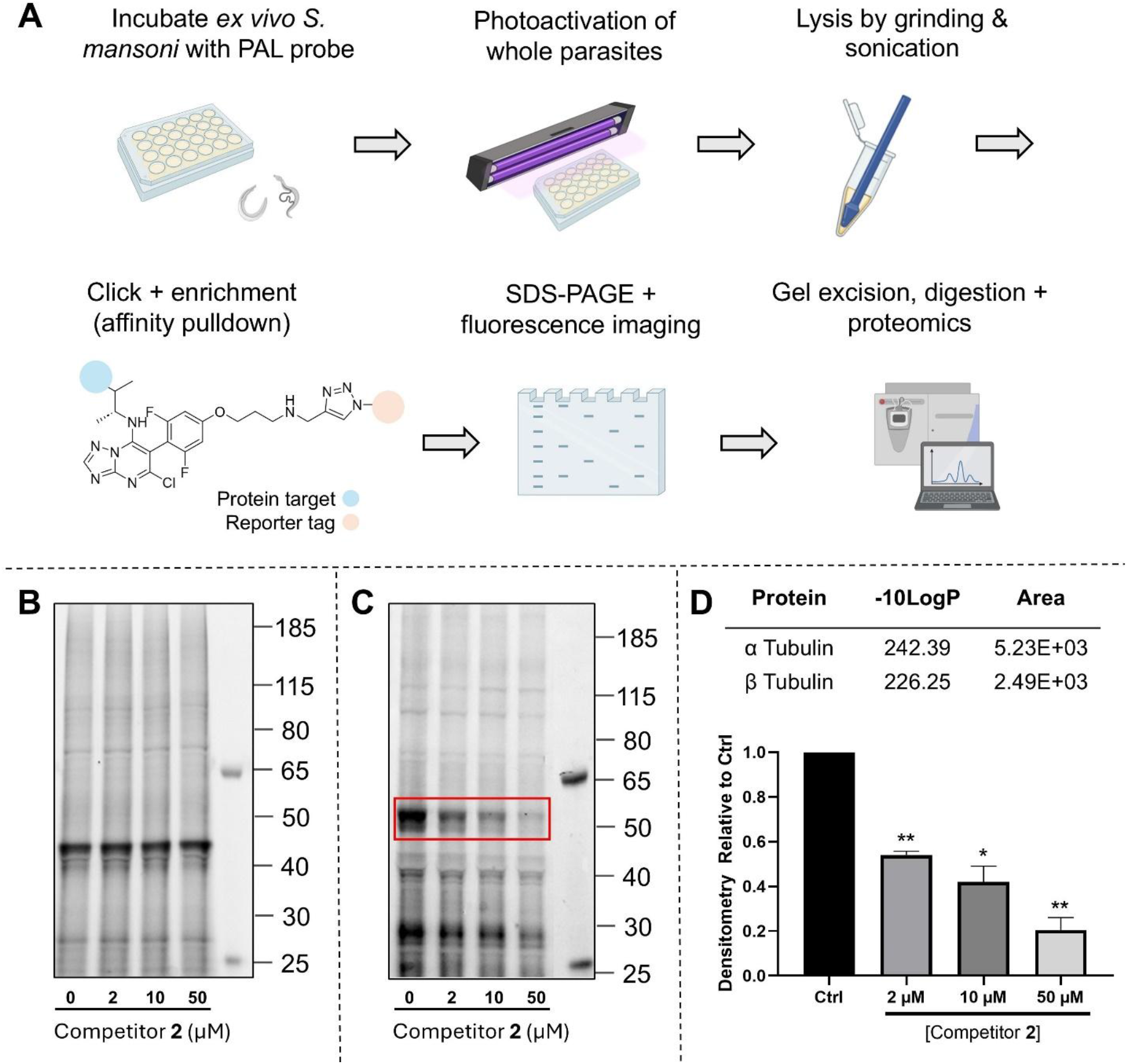
(**A**) Summary of the PAL protocol for *ex vivo S. mansoni* adults. (**B**) PAL experiments with *S. mansoni* lysates and (**C**) living worms resolve different labeling profiles under the same conditions with probe **1**. (**D**) Proteomics and densitometry analysis. Proteomics analysis of the protein band labeled specifically by **1** (red box in **C**) identifies *S. mansoni* tubulin. Densitometry analysis reveals concentration-dependent binding in presence of the photostable competitor, **2**. Protein concentrations were measured using the BCA assay and normalized to 0.3 µg/mL to ensure equal protein loading. *p < 0.005 and **p < 0.005 by one-way ANOVA.

To evaluate the specificity of binding by probe **1**, competition experiments were performed by pre-incubating worms with the photostable competitor, **2** (2, 10 and 50 µM), for 30 min prior to incubation with 2 µM **1**. The labeling workflow was also applied to *S. mansoni* lysates (see Supporting Information) to understand the probe’s binding specificity under such conditions, and if determined, whether binding could be competed for with the competitor.

With worm lysates, several fluorescent bands were observed by PAL with probe **1**, including a prominent band at ∼45 kDa (**Figure 2B**). The intensity of labeling of these bands could not be diminished in the presence of increasing concentrations of the competitor, **2**, suggesting non-specific binding. PAL performed with live worms also revealed several intense bands, with the most prominent at ∼55 kDa (**Figure 2C**). In the presence of increasing concentrations of competitor **2**, the fluorescence intensity of this band decreased markedly, indicating a specific competitive interaction (**Figure 2C, D**). Other bands, particularly at ∼40 and ∼28 kDa, were also labeled by probe **1**; however, their competition profiles were less pronounced than that for the 55 kDa band, suggestive of non-specific binding. After lysis of the live worms that had been exposed to PAL, the proteins bound by probe **1** were enriched by neutravidin-affinity purification. The 55 kDa gel band was excised and subjected to proteomic analysis, which identified the sample to be primarily composed of α- and β-tubulin (**Figure 2D**).

Collectively, whereas, PAL with worm lysates using **1** in competition with **2** suggested non-specific binding, the data generated with living worms confirms a specific PAL reaction with tubulin at 55 kDa. In the former experiment, it is possible that MTs depolymerize during lysate preparation, resulting in the loss of the vinca and seventh binding sites at the α-β tubulin interface. If confirmed, then the finding underscores the importance of investigating target binding under native conditions.

The demonstration here of a specific PAL reaction is, to the best of our knowledge, novel for a metazoan, and underscores the utility of this technique to identify potential drug targets from phenotypic screens which are commonly employed for this parasite. Importantly, with the associated knowledge of a small molecule’s chemistry and structure-activity relationship (SAR), nearly any bioactive ligand can potentially be adapted as a PAL probe, thereby opening opportunities to investigate a broad range of protein targets. In addition to target identification, PAL can be integrated with chemoproteomic approaches to enable a quantitative analysis of probe-bound proteins. We will apply this strategy in the future to quantify target-engagement and -selectivity. Overall, the results reported here highlight the potential of PAL as a versatile platform to support target-based drug discovery in the schistosome and evaluate protein-ligand interactions in other metazoans.

## ACKNOWLEDGEMENTS

The research reported was supported by the NIH grant R21AI156554 to CRC and CB. KRF acknowledges support of the CARING T32 Training Grant (T32AI007036).

## Supporting Information

*S. mansoni assays*. Maintenance of the *S. mansoni* life cycle (NMRI isolate), the perfusion of adult worms (≥42 days old) and the incubation with test compounds were as described.^41, 46^ The use of mice to maintain the *S. mansoni* life-cycle was approved by the Institutional Animal Care and Use Committee of the University of California San Diego. Parasite phenotypic responses were visually assessed at 72 h using a standardized set of descriptors, which were then converted to severity scores ranging from 0 (no activity) to 4 (maximum activity).^33, 41, 42^

*In silico studies*. Molecular modeling was carried out on a workstation (Micro-Star International Co., Ltd. MS-7D25, 12th Gen Intel® Core™ i9-12900 K, 24 cores) running Ubuntu 22.04.3 LTS and equipped with an NVIDIA RTX A5000 graphics card. The crystal structure of mammalian α- and β-tubulin was obtained from the Protein Data Bank (PDB ID: 7CLD^38^; http://www.rcsb.org/). The corresponding *S. mansoni* FASTA sequences (α-tubulin: UniProt ID Q26444; β-tubulin: UniProt ID A0A5K4F3J5) share 87 and 66% sequence identity, respectively. Homology models of *S. mansoni* α- and β-tubulin were generated with the Homology Model Builder in the Schrödinger Suite [Schrödinger Release 2024-3: BioLuminate, Schrödinger, LLC, New York, NY, 2024], using the mammalian α- and β-tubulin structures as templates. Model refinement was performed with MacroModel^47^ using the OPLS4 force field. Protein structures were then processed with the Protein Preparation Wizard.^48^ All co-crystallized molecules were removed except for the ligand and GDP. Protonation states of protein residues were adjusted to pH 7.0. Docking simulations were carried out with Glide.^49^ The grid was centered on the co-crystallized ligand, with the inner box set to 10 Å and the outer box to 20 Å. Ten docking poses were generated for both the mammalian and *S. mansoni* tubulin structures. The best-ranked poses were further refined and rescored using MM-GBSA.^50^ Pictures were prepared using PyMOL.^51^

### Photoaffinity Labeling

#### PAL with living adult S. mansoni

PAL experiments were performed in a biosafety cabinet using sterile techniques. In a 24-well microplate, five male and two female worms were incubated with 2 µM probe **1** (0.1% DMSO final) at 37 °C and 5% CO_2_ for 2 h in Basch medium^52^ containing 2.5% heat-inactvated FBS and 1X penicilin-streptomycin solution. For competition experiments, worms were first preincubated for 30 min with 50 µM of the photostable competitor, **2**, followed by addition of 2 µL PAL probe **1**. The microplate plates lids were removed, and the worms exposed for 30 min to a hand-held 365 nm light source positioned at a distance of 1 – 2 cm from the surface of the medium (no assay plate lid).^35^ After photoactivation, the medium was removed and the parasites were washed three times in PBS before freezing at -80 °C.

#### Homogenization

Worms were lysed in 200 µL PBS containing 1X protease inhibitor cocktail (without EDTA; Thermo Cat#1862209), using a Fisherbrand Model 120 Sonic Dismembrator probe sonicator. Pulses were for 2 sec with 5 sec pauses over a total of 10 min. The protein concentration was quantified using the Pierce™ BCA Protein Assay Kit (ThermoFisher Cat#23227) and samples normalized by diluting in PBS (0.3 mg/mL).

#### PAL with S. mansoni lysates

Untreated *S. mansoni* worms were homogenized as described above. In Eppendorf tubes, 200 µL of cell lysates (0.3 mg/mL) were incubated with 2 µM probe **1** (0.1% DMSO final) at 37 °C for 2 h. For competition experiments, lysates were pre-incubated for 30 min with 50 µM of the photostable competitor, **2**, followed by addition of 2 µL probe **1**. The resulting lysates were transferred to 96-well polypropylene plates and exposed for 30 min to a hand-held 365 nm light source positioned at a distance of 1 – 2 cm from the surface of lysate solution (no assay plate lid).

#### Click reaction

The Click reaction was conducted as previously reported,^35^ using 200 µL protein sample, 5 µL of a 5 mM solution of Biotin-TAMRA azide (Click Chemistry Tools Cat # 1048) and 25 µL Click catalyst mixture prepared in a 3:1:1 ratio of 1.7 mM Tris(benzyltriazolylmethyl)amine^53^ (TBTA; in 80% *t*-butanol and 20% DMSO) : 50 mM CuSO_4_ (in water) : 50 mM Tris (2-carboxyethyl) phosphine (TCEP; adjusted to pH 7.0 in water).^44^ The reaction mixture was then incubated at 37 °C for 45 min.

#### SDS-PAGE and fluorescence visualization

Clicked protein samples (30 µL) were added to 10 µL 4X LDS sample loading buffer (Fisher Cat# NP0007) containing 10% 2-mercaptoethanol (TCI Cat# M1948). The samples were then denatured at 95 °C for 2 min and loaded onto a precast Bolt™ 1.00 mm, 4-12% Bis-Tris plus mini-gel. Electrophoresis employed a constant 120V for 1.5 h. Protein bands were then visualized by fluorescence imaging using a Bio-Rad ChemiDoc MP imaging system and stained with Coomassie Brilliant Blue G250. Fluorescence signal intensities for the probe-competition experiments were densitometrically quantified using a Bio-Rad Image Lab 6.1.

#### Affinity purification

After the Click reaction, the protein mixture was precipitated by the addition of three volumes of cold acetone and incubated in a -20 °C freezer for 30 min. The precipitated protein was centrifuged at 2,000 *g* and the supernatant aspirated. The pellet was washed with cold acetone, resuspended, and centrifuged twice more. After the final acetone wash, the pellet was air dried and reconstituted in 1% SDS in PBS. The proteins in the pellet were subjected to affinity purification as previously described using 50% Neutravidin agarose beads (25 µL; Fisher Cat # 29200).^35^ The purified proteins were separated via SDS-PAGE, visualized by fluorescence, and stained with Coomassie Brilliant Blue G250, as described above. The gel bands of interest were excised for proteomics analysis.

#### In-gel digest and proteomics analysis

In-gel digestion of gel excisions was performed as previously reported.^35^ The resulting extracts were analyzed directly by ultra-high pressure liquid chromatography (UPLC) coupled with MS/MS using nano-spray ionization. The nanospray ionization experiments were performed using an Orbitrap Fusion Lumos Hybrid Mass Spectrometer (Thermo), interfaced with a nano-scale reversed-phase UPLC (Thermo Dionex UltiMate™ 3000 RSLC nano System) and a 25 cm, 75-µm ID glass capillary packed with 1.7-μm C18 (130) BEHTM beads (Waters), as previously reported.^35^ Protein identification was carried out using the Peaks Studio X software (Bioinformatics solutions Inc.).^54^

## References

1. McManus DP, Dunne DW, Sacko M, Utzinger J, Vennervald BJ, Zhou XN. Schistosomiasis. Nat Rev Dis Primers. 2018;4(1): 13.

2. Buonfrate D, Ferrari TCA, Adegnika AA, Russell Stothard J, Gobbi FG. Human schistosomiasis. Lancet. 2025;405(10479): 658–670.

3. Schistosomiasis; 2023. https://www.who.int/news-room/fact-sheets/detail/schistosomiasis.

4. Rinaldo D, Perez-Saez J, Vounatsou P, Utzinger J, Arcand JL. The economic impact of schistosomiasis. Infect Dis Poverty. 2021;10(1): 134.

5. Doruska MJ, Barrett CB, Rohr JR. Modeling how and why aquatic vegetation removal can free rural households from poverty-disease traps. Proc Natl Acad Sci U S A. 2024;121(52): e2411838121.

6. Andrews P, Thomas H, Pohlke R, Seubert J. Praziquantel. Med Res Rev. 1983;3(2): 147–200.

7. Waechtler A, Cezanne B, Maillard D, et al. Praziquantel - 50 Years of Research. ChemMedChem. 2023;18(12): e202300154.

8. Sisay M, Hailu T, Damtie D, Geta K, Zelalem L, Misganaw D. Efficacy of single dose praziquantel against Schistosoma mansoni in East Africa: a systematic review and meta-analysis. Sci Rep. 2025;15(1): 4642.

9. Hailegebriel T, Nibret E, Munshea A. Efficacy of Praziquantel for the Treatment of Human Schistosomiasis in Ethiopia: A Systematic Review and Meta-Analysis. J Trop Med. 2021;2021: 2625255.

10. Sabah AA, Fletcher C, Webbe G, Doenhoff MJ. Schistosoma mansoni: chemotherapy of infections of different ages. Exp Parasitol. 1986;61(3): 294–303.

11. Gönnert R, Andrews P. Praziquantel, a new board-spectrum antischistosomal agent. Z Parasitenkd. 1977;52(2): 129–150.

12. Bustinduy AL, Waterhouse D, de Sousa-Figueiredo JC, et al. Population Pharmacokinetics and Pharmacodynamics of Praziquantel in Ugandan Children with Intestinal Schistosomiasis: Higher Dosages Are Required for Maximal Efficacy. mBio. 2016;7(4).

13. Andrews P. Praziquantel: mechanisms of anti-schistosomal activity. Pharmacol Ther. 1985;29(1): 129–156.

14. Eastham G, Fausnacht D, Becker MH, Gillen A, Moore W. Praziquantel resistance in schistosomes: a brief report. Front Parasitol. 2024;3: 1471451.

15. Geary TG, Sakanari JA, Caffrey CR. Anthelmintic drug discovery: into the future. J Parasitol. 2015;101(2): 125–133.

16. Caffrey CR. Chemotherapy of schistosomiasis: present and future. Curr Opin Chem Biol. 2007;11(4): 433–439.

17. Caffrey C, El-Sakkary N, Mäder P, et al. Drug Discovery and Development for Schistosomiasis. Neglected Tropical Diseases: Drug Discovery and Development. 2019;77: 187–225.

18. Kalinna BH, Ross AG, Walduck AK. Schistosome Transgenesis: The Long Road to Success. Biology (Basel). 2024;13(1).

19. Li T, Yang Y, Qi H, et al. CRISPR/Cas9 therapeutics: progress and prospects. Signal Transduct Target Ther. 2023;8(1): 36.

20. Setten RL, Rossi JJ, Han SP. The current state and future directions of RNAi-based therapeutics. Nat Rev Drug Discov. 2019;18(6): 421–446.

21. Lye LF, Dobson DE, Beverley SM, Tung MC. RNA interference in protozoan parasites and its application. J Microbiol Immunol Infect. 2025;58(3): 281–287.

22. Pal S, Dam S. CRISPR-Cas9: Taming protozoan parasites with bacterial scissor. J Parasit Dis. 2022;46(4): 1204–1212.

23. Nance J, Frøkjær-Jensen C. The Caenorhabditis elegans Transgenic Toolbox. Genetics. 2019;212(4): 959–990.

24. Jurberg AD, Brindley PJ. Gene function in schistosomes: recent advances toward a cure. Front Genet. 2015;6: 144.

25. Ittiprasert W, Brindley PJ. CRISPR-based functional genomics for schistosomes and related flatworms. Trends Parasitol. 2024;40(11): 1016–1028.

26. Quinzo MJ, Perteguer MJ, Brindley PJ, Loukas A, Sotillo J. Transgenesis in parasitic helminths: a brief history and prospects for the future. Parasit Vectors. 2022;15(1): 110.

27. Skelly PJ, Da’dara A, Harn DA. Suppression of cathepsin B expression in Schistosoma mansoni by RNA interference. Int J Parasitol. 2003;33(4): 363–369.

28. Boyle JP, Wu XJ, Shoemaker CB, Yoshino TP. Using RNA interference to manipulate endogenous gene expression in Schistosoma mansoni sporocysts. Mol Biochem Parasitol. 2003;128(2): 205–215.

29. Li J, Xiang M, Zhang R, Xu B, Hu W. RNA interference in vivo in Schistosoma japonicum: Establishing and optimization of RNAi mediated suppression of gene expression by long dsRNA in the intra-mammalian life stages of worms. Biochem Biophys Res Commun. 2018;503(2): 1004–1010.

30. Wang J, Paz C, Padalino G, et al. Large-scale RNAi screening uncovers therapeutic targets in the parasite. Science. 2020;369(6511): 1649–1653.

31. Stephens DR, Fung HYJ, Han Y, et al. A genome-scale drug discovery pipeline uncovers therapeutic targets and a unique p97 allosteric binding site in. Proc Natl Acad Sci U S A. 2025;122(35): e2505710122.

32. Xie Y, Wang X, Cheng S, et al. RNAi screening of uncharacterized genes identifies promising druggable targets in Schistosoma japonicum. PLoS Pathog. 2025;21(3): e1013014.

33. Monti L, Cornec AS, Oukoloff K, et al. Congeners Derived from Microtubule-Active Phenylpyrimidines Produce a Potent and Long-Lasting Paralysis of Schistosoma mansoni In Vitro. ACS Infect Dis. 2021;7(5): 1089–1103.

34. Monti L, Liu LJ, Varricchio C, et al. Structure-Activity Relationships, Tolerability and Efficacy of Microtubule-Active 1,2,4-Triazolo[1,5-a]pyrimidines as Potential Candidates to Treat Human African Trypanosomiasis. ChemMedChem. 2023;18(20): e202300193.

35. Lucero B, Francisco KR, Varricchio C, et al. Design, Synthesis, and Evaluation of An Anti-trypanosomal [1,2,4]Triazolo[1,5-a]pyrimidine Probe for Photoaffinity Labeling Studies. ChemMedChem. 2024;19(8): e202300656.

36. Oukoloff K, Kovalevich J, Cornec AS, et al. Design, synthesis and evaluation of photoactivatable derivatives of microtubule (MT)-active [1,2,4]triazolo[1,5-a]pyrimidines. Bioorg Med Chem Lett. 2018;28(12): 2180–2183.

37. Alle T, Varricchio C, Yao Y, et al. Microtubule-Stabilizing 1,2,4-Triazolo[1,5-a]pyrimidines as Candidate Therapeutics for Neurodegenerative Disease: Matched Molecular Pair Analyses and Computational Studies Reveal New Structure–Activity Insights. J Med Chem. 2023;66(1): 435–459.

38. Sáez-Calvo G, Sharma A, Balaguer FA, et al. Triazolopyrimidines Are Microtubule-Stabilizing Agents that Bind the Vinca Inhibitor Site of Tubulin. Cell Chem Biol. 2017;24(6): 737-750.e736.

39. Yang J, Yu Y, Li Y, et al. Cevipabulin-tubulin complex reveals a novel agent binding site on α-tubulin with tubulin degradation effect. Sci Adv. 2021;7(21).

40. Kovalevich J, Cornec AS, Yao Y, et al. Characterization of Brain-Penetrant Pyrimidine-Containing Molecules with Differential Microtubule-Stabilizing Activities Developed as Potential Therapeutic Agents for Alzheimer’s Disease and Related Tauopathies. J Pharmacol Exp Ther. 2016;357(2): 432–450.

41. Abdulla MH, Ruelas DS, Wolff B, et al. Drug discovery for schistosomiasis: hit and lead compounds identified in a library of known drugs by medium-throughput phenotypic screening. PLoS Negl Trop Dis. 2009;3(7): e478.

42. Long T, Rojo-Arreola L, Shi D, et al. Phenotypic, chemical and functional characterization of cyclic nucleotide phosphodiesterase 4 (PDE4) as a potential anthelmintic drug target. PLoS Negl Trop Dis. 2017;11(7): e0005680.

43. Francisco KR, Monti L, Yang W, et al. Structure-activity relationship of dibenzylideneacetone analogs against the neglected disease pathogen, Trypanosoma brucei. Bioorg Med Chem Lett. 2023;81: 129123.

44. Mackinnon AL, Taunton J. Target Identification by Diazirine Photo-Cross-linking and Click Chemistry. Curr Protoc Chem Biol. 2009;1: 55–73.

45. Kolb H, Finn M, Sharpless K. Click chemistry: Diverse chemical function from a few good reactions. Angewandte Chemie-International Edition. 2001;40(11): 2004-+.

46. Abdulla MH, Lim KC, Sajid M, McKerrow JH, Caffrey CR. Schistosomiasis mansoni: novel chemotherapy using a cysteine protease inhibitor. PLoS Med. 2007;4(1): e14.

47. Schrödinger Release 2024-3: MacroModel. New York, NY: Maestro, Schrödinger, LLC; 2024.

48. Schrödinger Release 2024-3: Protein Preparation Workflow; Epik. New York, NY: Maestro, Schrödinger, LLC; 2024.

49. Schrödinger Release 2024-3: Glide. New York, NY: Maestro, Schrödinger, LLC; 2024.

50. Schrödinger Release 2024-3: Prime. New York, NY: Maestro, Schrödinger, LLC; 2024.

51. The PyMOL Molecular Graphics System, Version 2.0 Schrödinger, LLC.

52. Basch PF. Cultivation of Schistosoma mansoni in vitro. I. Establishment of cultures from cercariae and development until pairing. J Parasitol. 1981;67(2): 179–185.

53. Chan TR, Hilgraf R, Sharpless KB, Fokin VV. Polytriazoles as copper(I)-stabilizing ligands in catalysis. Org Lett. 2004;6(17): 2853–2855.

54. Zhang J, Xin L, Shan B, et al. PEAKS DB: de novo sequencing assisted database search for sensitive and accurate peptide identification. Mol Cell Proteomics. 2012;11(4): M111.010587.

